## Supplemental Figures for "Longitudinal development of category representations in ventral temporal cortex predicts word and face recognition"

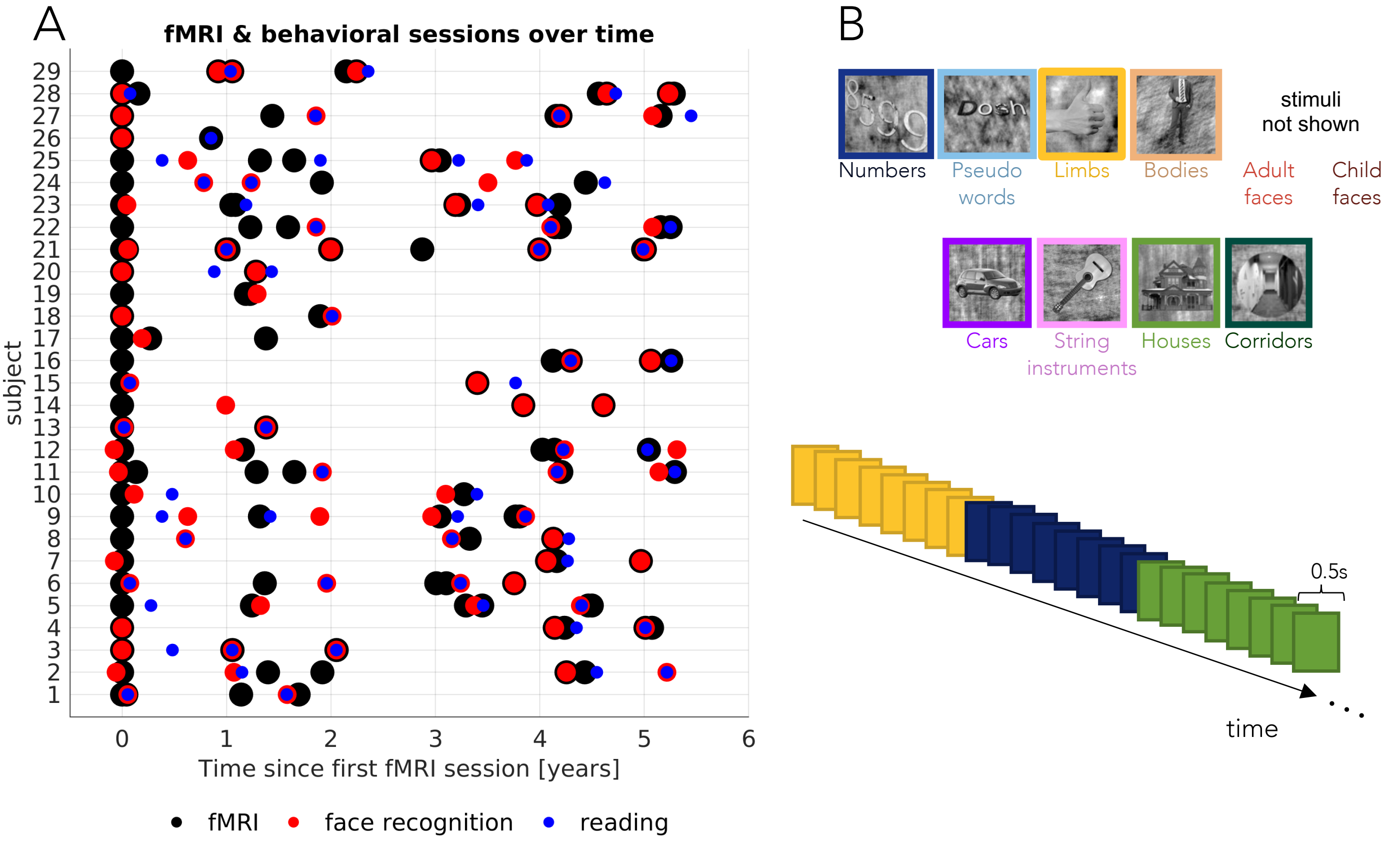


**Fig S1. Data acquisition and experimental design**

(A) fMRI and behavioral sessions in each participant over time**.** The scatter plot illustrates the number and temporal sequence of fMRI and behavioral sessions in each child included in the analyses. Each row is a participant. Sessions are plotted relative to each child's first fMRI session. (B) Example stimuli and fMRI experimental design. *Top:* example stimuli for each of the 10 categories, which can be grouped into five domains (characters, body parts, faces, objects, and places). *Bottom:* Schematic illustrating the fMRI paradigm. Stimuli were shown at a rate of 2Hz and were presented in random order 4s blocks. Participants were instructed to fixate on a central dot and indicate with a button press when an image that only contained a scrambled background appeared.

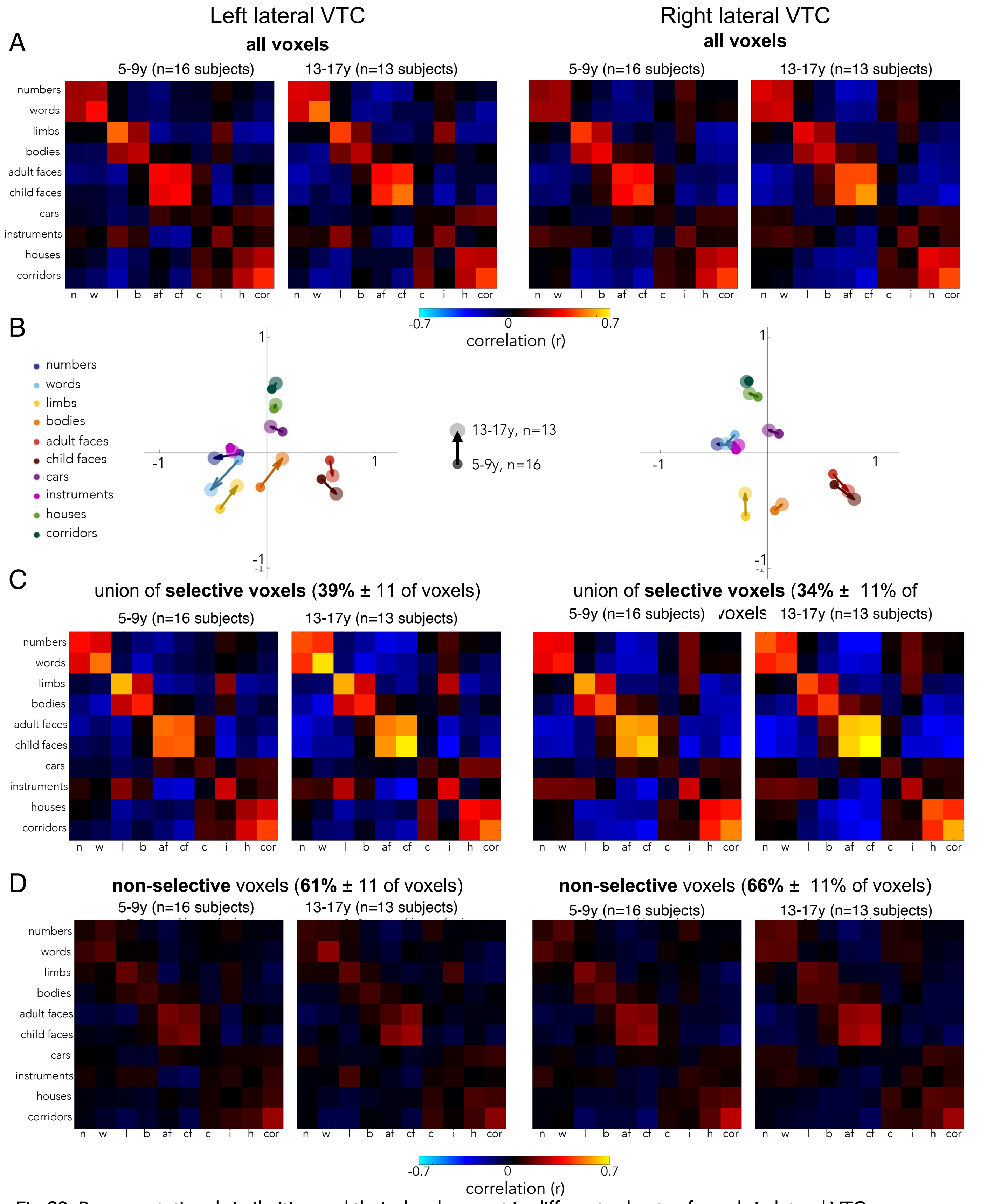


**Fig S2. Representational similarities and their development in different subsets of voxels in lateral VTC.**
(A) Representational similarity matrices (RSM) of left and right lateral VTC for 5-9-year-olds and 13-17-year-olds. One session per child is included per RSM of each age group. (B) Multidimensional scaling (MDS) embeddings for the category representation for the same age groups: 5-9year-olds (small circles) and 13-17-year-olds (larger circles). (C,D) Same as (A) but for different subsets of voxels; the union of the selective voxels (C) and the non-selective voxels (D). The color scale is the same across (A,C,D).

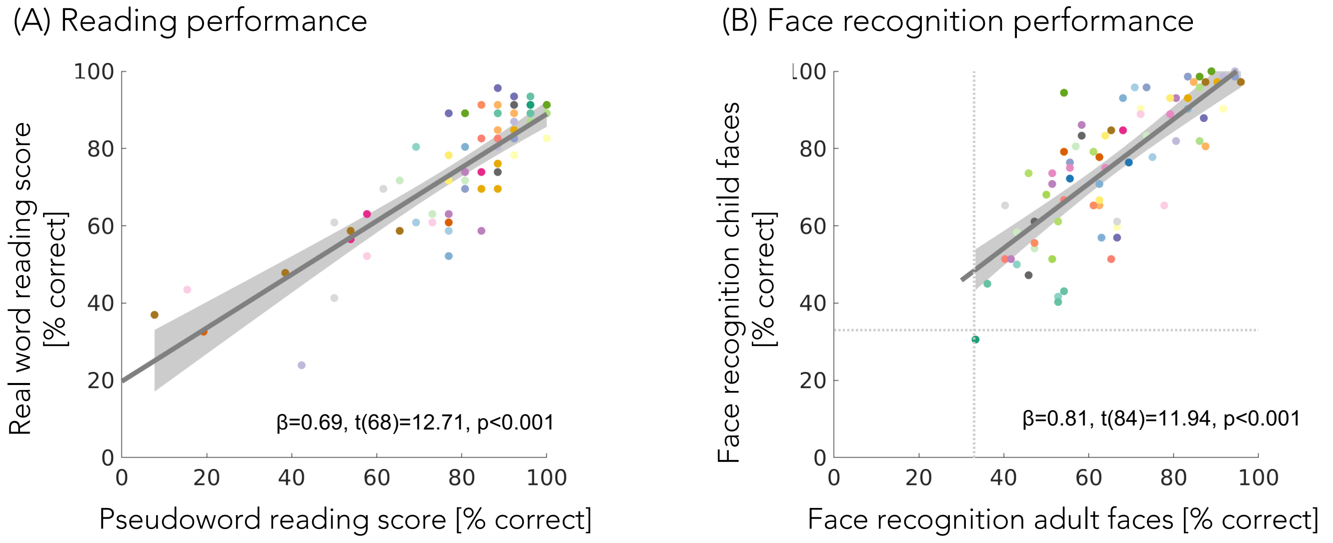


**Fig S3. Reading and face recognition performance are reliable across tests**
(A) Reading performance on pseudowords (x-axis) predicts reading performance on real words (y-axis). (B) Face recognition performance in the Cambridge face recognition task (CFMT) with images of adults faces (x-axis) predicts performance on the CFMT test with images of child faces (y-axis). Dashed line indicates chance level in the CFMT. Subjects are coded by color. *Gray line:* Linear mixed model (LMM) prediction. *Shaded gray:* 95% confidence interval (CI)
